# LxxIxE-like Motif in Spike Protein of SARS-CoV-2 that is Known to Recruit the Host PP2A-B56 Phosphatase Mimics Artepillin C, an Immunomodulator, of Brazilian Green Propolis

**DOI:** 10.1101/2020.04.01.020941

**Authors:** Halim Maaroufi

## Abstract

SARS-CoV-2 is highly contagious and can cause acute respiratory distress syndrome (ARDS) and multiple organ failure that are largely attributed to the cytokine storm. The surface coronavirus spike (S) glycoprotein is considered as a key factor in host specificity because it mediates infection by receptor-recognition and membrane fusion. Here, the analysis of SARS-CoV-2 S protein revealed two B56-binding LxxIxE-like motifs in S1 and S2 subunits that could recruit the host protein phosphatase 2A (PP2A). The motif in S1 subunit is absent in SARS-CoV and MERS-CoV. Phosphatases and kinases are major players in the regulation of pro-inflammatory responses during pathogenic infections. Moreover, studies have shown that viruses target PP2A in order to manipulate host’s antiviral responses. Recent researches have indicated that SARS-CoV-2 is involved in sustained host inflammation. Therefore, by controlling acute inflammation, it is possible to eliminate its dangerous effects on the host. Among efforts to fight COVID-19, the interaction between LxxIxE-like motif and the PP2A-B56-binding pocket could be a target for the discovery and/or development of a bioactive ligand inhibitor for therapeutic purposes. Indeed, a small molecule called Artepillin C (ArtC), a main compound in Brazilian honeybee green propolis, mimics the side chains of LxxLxE motif. Importantly, ArtC is known, among other effects, to have anti-inflammatory activity that makes it an excellent candidate for future clinical trials in COVID-19 patients.

## INTRODUCTION

In March 11^th^ 2020, the World Health Organization (WHO) announced that COVID-19 (<u>Co</u>rona<u>vi</u>rus <u>D</u>isease-20<u>19</u>) situation is a pandemic. The novel SARS-CoV-2 has had serious consequences for human health and socioeconomic stability worldwide. Coronaviruses (CoVs) are a large family of enveloped single positive-stranded RNA viruses that can infect both mammalian and avian species because their rapid mutation and recombination facilitate their adaptation to new hosts (Graham and Baric, 2010; Li, 2013). They can cause severe, often fatal Acute Respiratory Distress Syndrome (ARDS). CoVs are classified into *Alpha-, Beta-, Gamma-*, and *Deltacoronavirus* genetic genera. The novel betacoronavirus (betaCoVs) SARS-CoV-2 is relatively close to other betaCoVs: severe acute respiratory syndrome coronavirus (SARS-CoV), Middle East respiratory syndrome coronavirus (MERS-CoV), bat coronavirus HKU4, mouse hepatitis coronavirus (MHV), bovine coronavirus (BCoV), and human OC43 coronavirus (HCoV-OC43). SARS-CoV emerged in China (2002–2003) and spread to other countries (more than 8,000 infection cases and a fatality rate of ~10%) (Peiris et al., 2003). In 2012, MERS-CoV was detected in the Middle East. It spread to multiple countries, infecting more than 1,700 people with a fatality rate of ~36% (de Wit et al., 2016).

The surface-located SARS-CoV-2 spike glycoprotein S (S) is a 1273 amino acid residues. It is a homotrimeric, multidomain, and integral membrane protein that give coronaviruses the appearance of having crowns (*Corona* in Latin) (Li, 2016). It is a key piece of viral host recognition (receptor-recognition) and organ tropism and induces strongly the host immune reaction (Li, 2015). It is subdivided to S1 subunit that binds to a receptor on the host cell surface and S2 subunit that permits viral and host membranes fusion. S1 subunit is divided into two domains, an N-terminal domain (NTD) and a C-terminal receptor-binding domain (RBD) that can function as viral receptors-binding (Li, 2012). In addition, S1 subunit is normally more variable in sequence among different CoVs than is the S2 subunit (Masters, 2006).

Protein phosphatase 2A (PP2A) is a major family of Serine/Threonine phosphatases in eukaryotic cells and regulates diverse biological processes through dephosphorylation of numerous signaling molecules. PPA2 and phosphatase 1 (PP1), regulates over 90% of all Ser/Thr dephosphorylation events in eukaryotic cells (Eichhorn et al., 2009). PP2A is a heterotrimeric holoenzyme composed of a stable heterodimer of the scaffold A-subunit (PP2A-A) and catalytic C-subunit (PP2A-C) and a variable mutually exclusive regulatory subunit from four families (B (B55), B’ (B56), B’’ and B’’’) which provide substrate specificity. The human B56 family consists of at least five different members (α, β, γ, δ and ε). Phosphatases and kinases are big players in the regulation of pro-inflammatory responses during microbial infections. Sun et al. (2017) showed that PP2A plays an important role in regulating inflammation by controlling the production of inflammatory cytokines/chemokines (Kozicky and Sly, 2015). In addition, PP2A is one of the phosphatases involved in negatively regulating the inflammatory response (Shanley et al., 2001). Moreover, studies have revealed that viruses use multiple strategies to target PP2A in the aim to manipulate host antiviral responses (Guergnon et al., 2011).

Artepillin C (ArtC) is uniquely found in Brazilian honeybee green propolis and is one of its major bioactive components (Marcucci et al., 2001; Park et al., 2004). It is a low-molecular weight phenolic single ring with two prenyl groups (3,5-diprenyl-4-hydroxycinnamic acid) (Szliszka et al., 2013). These properties suggest high oral bioavailability and cell-permeability allowing good biological activity of ArtC (Shimizu et al., 2004; Konishi et al. 2005; Konishi, 2005). Indeed, Paulino et al. (2008) showed that ArtC exhibited bioavailability by oral administration in mice. Interestingly, ArtC has many therapeutic effects, anti-microbial, anti-tumor, apoptosis-inductor, immunomodulatory, and anti-oxidant effects (Salomão et al., 2004; Kimoto et al., 2001; Orsolic et al., 2006; Matsuno et al., 1997; Gekker et al., 2005; Nakanishi et al., 2003). Many of these therapeutic effects can be attributed to its immunomodulatory functions (Chan et al., 2013; Cheung et al., 2011; Paulino, et al., 2008). Indeed, Szliszka et al., (2013) have tested the anti-inflammatory activity of ArtC in activated RAW264.7 macrophages. They found that ArtC exerted strong antioxidant activity and significantly inhibited the production of several pro-inflammatory cytokines, such as TNF-α, IL-1β and IL-12, which makes ArtC an excellent anti-inflammatory drug. In addition, ArtC suppresses T cell proliferation and activation (Chan et al., 2013). Here, S protein was analyzed because of its importance in mediating infection. This analysis revealed two B56-binding LxxIxE-like motifs in S1 and S2 subunits that could recruit the host PP2A. Interestingly, side chains of LxxLxE motif present similarity with a small molecule called Artepillin C (ArtC), a main compound in Brazilian honeybee green propolis. Moreover, ArtC has anti-inflammatory activity that makes it an excellent candidate for future clinical trials in COVID-19 patients.

## RESULTS AND DISCUSSION

### Two LxxIxE-like motifs in S1 and S2 subunits of Spike protein

Sequence analysis of SARS-CoV-2 spike protein by the eukaryotic linear motif (ELM) resource (http://elm.eu.org/) revealed short linear motifs (SLiMs) known as LxxIxE-like motif ^293^**L**DP**L**S**E**^298^ in S1 subunit and ^1197^**L**ID**L**Q**E**^1202^ in S2 subunit (Fig. 1). SLiMs are few amino acid residues (3-15) in proteins that facilitate protein sequence modifications and protein-protein interactions (Davey et al., 2012; Van Roey et al., 2014). RNA viruses are known to mutate quickly and thus are able to create mimic motifs, on very short time scales, that could hijack biological processes in the host cell such as cell signaling networks (Davey et al., 2015; Via et al., 2015; Davey et al., 2011). Interestingly, ^293^LDPLSE^298^ is only present in SARS-CoV-2 and absent in S protein of coronaviruses analysed in this study (Fig. 1A). In order to interact with protein(s), ^293^LDPLSE^298^ must be present at the surface of S1 subunit. Indeed, this motif is exposed in the surface of S1 subunit in the end of NTD (Fig. 3B). A second motif ^1197^LIDLQE^1202^ is present in S2 subunit and conserved in SARS-CoV-2, SARS-CoV, SARS-like of bat from China and Kenya (Fig. 1B). These last betacoronaviruses are phylogenetically close (Fig. 2). Unfortunately, the region containing ^1197^LIDLQE^1202^ peptide has not been resolved in all known 3D structures of S protein to know if it is exposed in the surface. Probably, ^1197^LIDLQE^1202^ peptide is in an intrinsic disordered region of S protein.

**Figure 1.**
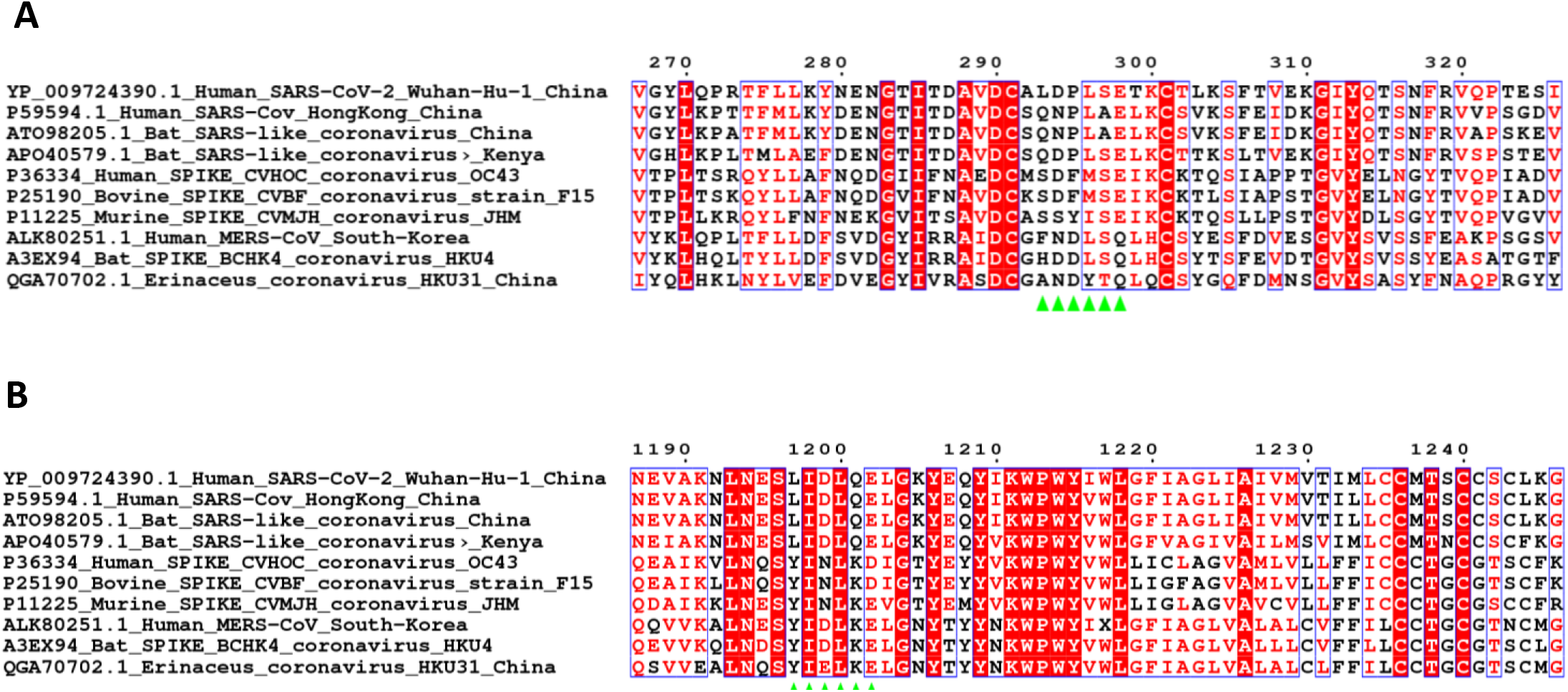
Multiple alignment of the spike glycoprotein of betacoronaviruses using Clustal omega. ^293^LDPLSE^298^ (A) and ^1197^LIDLQE^120^ (B) motifs are indicated by green stars. GenBank and UniProt accession numbers are indicated at the start of each sequence. The figure was prepared with ESPript (http://espript.ibcp.fr).

**Figure 2.**
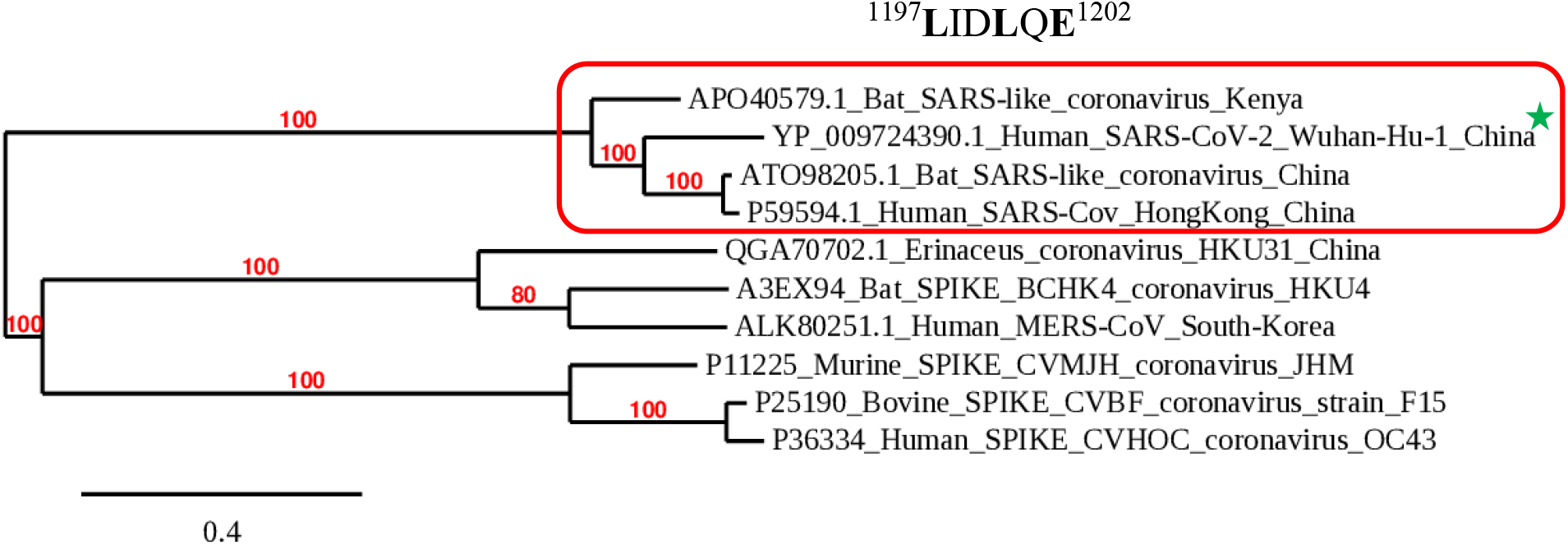
Unrooted phylogenetic tree of spike protein of representative betacoronaviruses. The tree was constructed using Mr Bayes method based on the multiple sequence alignment by Clustal omega. Red rectangle assembles betacoronaviruses with the same ^1197^LIDLQE^1202^. Green star indicated the only betacoronavirus with ^293^LDPLSE^298^. GenBank and UniProt accession numbers are indicated at the start of each sequence.

**Figure 3.**
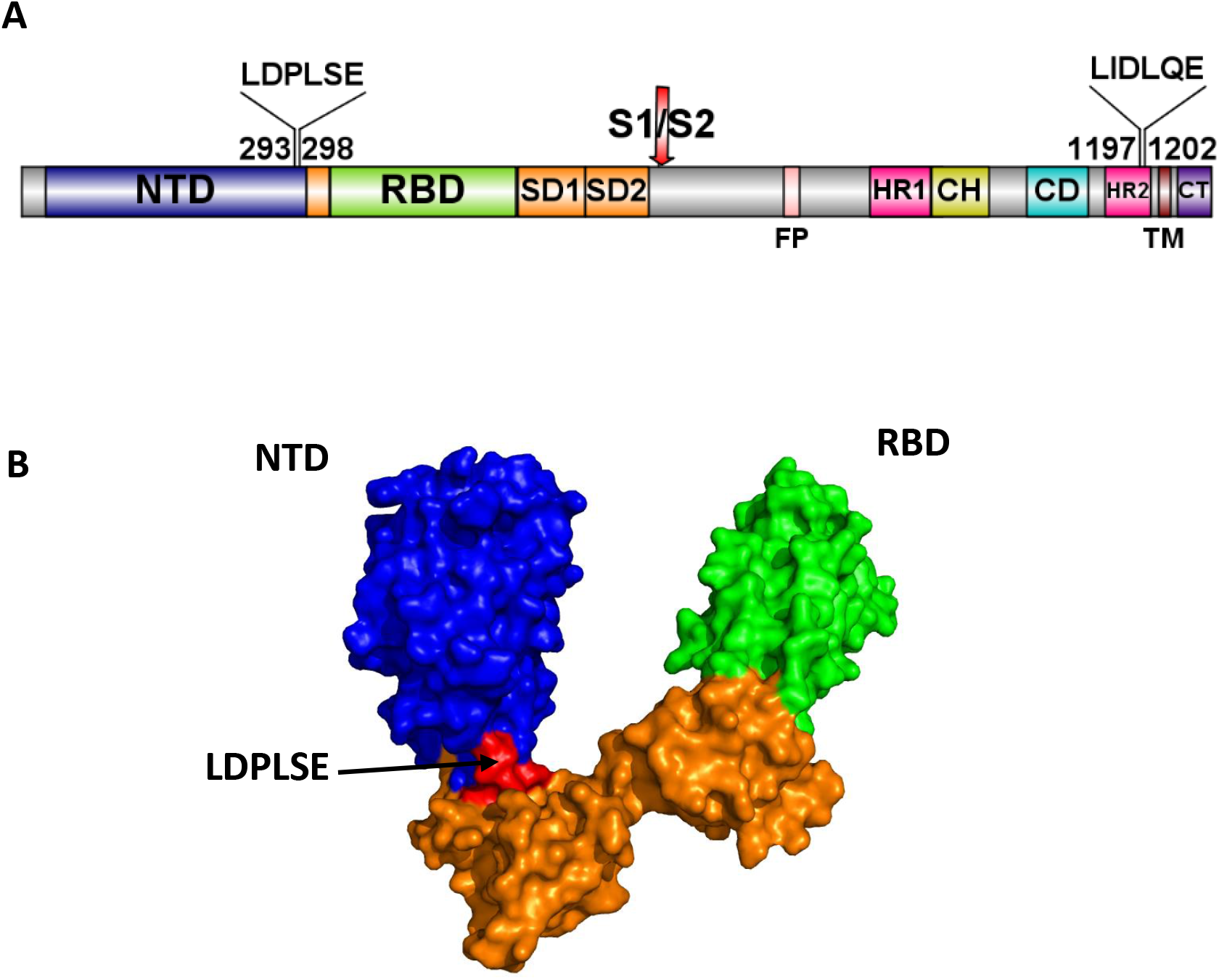
Spike (S) protein of SARS-CoV-2. **(A)** Diagram representation of S protein colored by domain. N-terminal domain (NTD), receptor-binding domain (RBD), subdomains 1 and 2 (SD1-2, orange), S1/S2 protease cleavage site, Fusion peptide (FP), heptad repeat 1 and 2 (HR1 and HR2), central helix (CH), connector domain (CD), transmembrane domain (TM), cytoplasmic tail (CT), and the localization of ^293^LDPLSE^298^ in the end of NTD and ^1197^LIDLQE^1202^ peptide in HR2. **(B)** Surface structure representation of the S1 subunit (PDBid: 6VSB_A). ^293^LDPLSE^298^ peptide is localized in the surface S1 subunit (red).

### Artepillin C, anti-inflammatory compound, mimics LxxLxE motif in S protein

In order to find small molecules with substituents that topologically and structurally resemble key amino acid side chains in LxxLxE motif, the method developed by Baran et al. (2007) has been used. This method allowed to discover a small molecule called Artepillin (ArtC) (Fig. 4A). It is a low-molecular weight phenolic single ring with two prenyl groups (3,5-diprenyl-4-hydroxycinnamic acid) (Szliszka et al., 2013). Figure 4B shows that the side chains of the two leucine and glutamic acid that constitute the key amino acid side chains are superimposed with the two prenyl groups and acid group of ArtC, respectively. ArtC is uniquely found in Brazilian honeybee green propolis and is one of its major bioactive components (Marcucci et al., 2001; Park et al., 2004). In addition, it has many therapeutic effects, anti-microbial, anti-tumor, apoptosis-inductor, immunomodulatory, and anti-oxidant effects (Salomão et al., 2004; Kimoto et al., 2001; Orsolic et al., 2006; Matsuno et al., 1997; Gekker et al., 2005; Nakanishi et al., 2003). Many of these therapeutic effects can be attributed to its immunomodulatory functions (Chan et al., 2013; Cheung et al., 2011; Paulino, et al., 2008).

**Figure 4.**
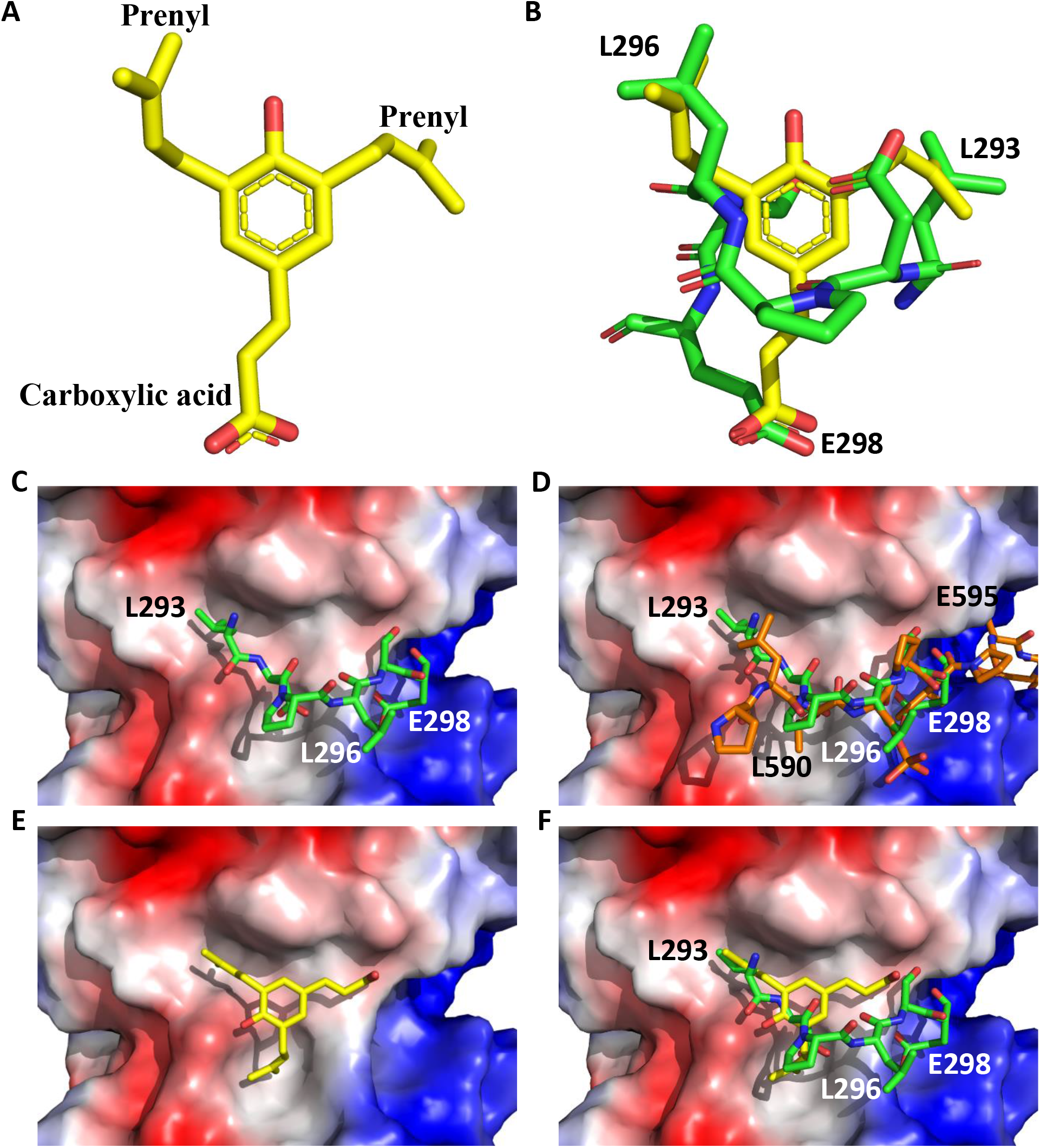
Stick representation of **(A)** Artepillin C (ArtC) and **(B)** superposition of ArtC (yellow) and ^293^LDPLSE^298^ peptide (green). Electrostatic potential surface representation of the region of the B56 regulatory subunit of PP2A (PDBid: 5SWF_A) with docked **(C)** ^293^LDPLSE^298^ peptide (green), **(D)** ^293^LDPLSE^298^ superimposed to pS-RepoMan (orange) (^581^rdiaskk<u>PL**Lp**SP**I**P**E**</u>LPEvpe^601^) peptide (PDBid: 5SW9_B), **(E)** ArtC (yellow) and **(F)** ArtC superimposed to ^293^LDPLSE^298^ peptide (green). The surfaces are colored by electrostatic potential with negative charge shown in red and positive charge in blue. Images were generated using PyMol (www.pymol.org).

### Interactions of ^293^LDPLSE^298^ and ArtC with B56 regulatory subunit

In order to determine the molecular interactions of ^293^LDPLSE^298^and ArtC with B56 regulatory subunit of PP2A (PP2A-B56), molecular docking was performed with the software AutoDock vina (Trott and Olson, 2010). Figure 4C and D show that ^293^LDPLSE^298^ peptide is localized in the same region as pS-RepoMan peptide that contains the LxxIxE motif (PDBid: 5SW9_B) and important amino acids of LxxIxE-like motif are superimposed with those of pS-RepoMan peptide (Fig. 4D). That is confirmed the reliability of AutoDock vina peptide docking module. In addition, Leu293 of ^293^LDPLSE^298^ is docked into hydrophobic pocket and Glu298 form ionic interactions with amino acid residues in positive charged region of PP2A-B56 (Fig. 4C). Note that ^293^LDPLSE^298^ contains a serine that could be phosphorylated generating a negative charge that will interact with positive patch in B56 subunit, enhancing binding affinity (Nygren and Scott, 2015). In the case of ArtC, its two prenyl groups are docked into two pockets as already seen with ^293^LDPLSE^298^ peptide (Fig. 4E-F). Note that the side chain of carboxylic acid group of ArtC is short to form ionic interactions with amino acid residues in positive charged region of PP2A-B56 (Fig. 4E). Therefore, to enhance binding affinity of ArtC, it is necessary to increase the length of side chain of carboxylic acid group. Interestingly, by using PinaColada, a computational method (Zaidman and Wolfson, 2016) for the design and affinity improvement of peptides that preclude protein-protein interaction, the peptide predicted (^293^LIDL<u>**E**</u>E^298^) by the software is mutated in C-terminal to negative charge residue (glutamic acid), showing the importance of the negative charge of peptide to interact with PP2A-B56. According to Autodock software, predicted binding affinity of ^293^LDPLSE^298^is −4.9 Kcal/mol, and this of ArtC is −6.1 Kcal/mol. It is known that the binding affinity of SLiMs is relatively weak (low μmolar range) (Gouw et al., 2018). This suggests that ArtC could compete with the virus to bind to PP2A-B56. To my knowledge, no compound has been identified to interact with regulatory subunits of PP2A, in addition it is the first time that a small molecule has been found that mimics the LxxIxE motif. Despite of numerous studies on ArtC its target is not yet known. In this study, its target is predicted as B56-PP2A. In general, activation of PP2A appears to have a suppressive effect on the inflammatory response (Sun et al., 2017). This suggests according to anti-inflammatory effect of ArtC that it could activate, *in vivo*, B56-PP2A.

### Protein phosphatase 2A and single RNA viruses

It has been shown in single RNA viruses, Ebola virus (EBOV) and Dengue fever virus (DENV) that they recruit the host PP2A through its regulatory subunit B56-binding LxxIxE motif to activate transcription and replication (Kruse et al., 2018; Oliveira et al., 2018). In addition, it has been shown an exacerbation of lung inflammation in mice infected with rhinovirus 1B (the most common viral infectious agent in humans). Administrating Salmeterol (beta-agonist) treatment to mice exerts anti-inflammatory effects by interacting with catalytic subunit PP2A, thus increasing its activity. It is probable that beta-agonists have the potential to target distinct pro-inflammatory pathways unresponsive to corticosteroids in patients with rhinovirus-induced exacerbations (Hatchwell et al., 2014). Treatment with Salmeterol drug may merit investigation for the possibility of using it in COVID-19’s patients with sustained and dangerous inflammatory reaction.

### Summary and conclusion

Analysis of SARS-CoV-2 proteome in the search of SLiMs that could be used by SARS-CoV-2 to manipulate host, allowed to discover in S protein an LxxIxE-like motif that is known to recruit the host PP2A-B56 phosphatase. Interestingly, PP2A is involved in the regulation of pro-inflammatory responses during pathogen infections. Well, recent researches have indicated that SARS-CoV-2 is involved in sustained host inflammation. Therefore, by controlling acute inflammation, it is possible to eliminate its dangerous effects on the host. LxxLxE motif of CoV-2 allowed to find a small molecule called Artepillin (ArtC), a main compound in Brazilian honeybee green propolis, which is known to have anti-inflammatory activity. ArtC, by its non-cytotoxicity in cells, high oral bioavailability, tested in mice, and cell-permeability, in addition that it can be synthesized in the laboratory (Uto et al., 2002; Yashiro et al., 2015) and produced in yeast by using synthetic biology (Munakata et al., 2019) makes it an ideal molecule for future clinical trials in COVID-19 patients.

## MATERIALS AND METHODS

### Sequence analysis

To search probable short linear motifs (SLiMs), SARS-CoV-2 spike protein sequence was scanned with the eukaryotic linear motif (ELM) resource (http://elm.eu.org/).

In the aim to find small molecules containing amino acids substituents that mimic LxxIxE-like motif, method described by Baran et al. (2007) was used.

### 3D modeling and molecular docking

3D structure of Artepillin C (ArtC) was obtained from PubChem database: https://pubchem.ncbi.nlm.nih.gov/compound/5472440#section=3D-Conformer.

For docking, the coordinates of the ^293^LDPLSE^298^ peptide were extracted from spike S protein of CoV-2 structure (PDBid: 6VSB_A). Unfortunately, the region containing ^1197^LIDLQE^1202^ peptide has not been resolved in all known 3D structures of spike S protein. So, Pep-Fold (Thevenet et al., 2012) software was used to model *de novo* this peptide. The model quality of the peptide was assessed by analysis of a Ramachandran plot through PROCHECK (Vaguine et al., 1999).

The docking of the two peptides into B56 regulatory subunit of PPA2 (PDBid: 5SWF_A) was performed with the software AutoDock vina (Trott and Olson, 2010). The 3D complex containing B56 subunit and peptides was refined by using FlexPepDock (London et al., 2011), which allows full flexibility to the peptide and side-chain flexibility to the receptor. The electrostatic potential surface of the B56 subunit was realized with PyMOL software (http://pymol.org/).

PinaColada a computational method (Zaidman and Wolfson, 2016) for inhibitory peptide design was used to improve affinity of ^293^LDPLSE^298^ peptide to bind to B56 regulatory subunit of PPA2. The software mutates several times the input peptide in the aim to find the highest binding affinity.

### Phylogeny

To establish the phylogenetic relationships between spike S protein of SARS-CoV-2 and representative betacoronaviruses, amino acid residues sequences were aligned with Clustal omega (Sievers et al., 2011) and a phylogenetic tree was constructed with MrBayes (Huelsenbeck and Ronquist, 2001) using: Likelihood model (Number of substitution types: 6(GTR); Substitution model: Poisson; Rates variation across sites: Invariable + gamma); Markov Chain Monte Carlo parameters (Number of generations: 100 000; Sample a tree every: 1000 generations) and Discard first 500 trees sampled (burnin).

## ACKNOWLEDGMENTS

I would like to thank the IBIS bioinformatics group for their help. I am grateful to Dr. Ahmad Abdel-Mawgoud SALEH, Université Laval, for the revision of the manuscript.

## CONFLICT OF INTERESTED

The author declares that he has no conflicts of interest.

## Notes

### Competing Interest Statement

The authors have declared no competing interest.

### Summary of Updates

-Addition of one section concerning Artepillin C, an Immunomodulator, of Brazilian Green Propolis that mimics LxxLxE motif (detailed description). -Modification of Figure 4.

